# Chromosomes distribute randomly to, but not within, human neutrophil nuclear lobes

**DOI:** 10.1101/2020.10.05.326009

**Authors:** Christine R Keenan, Michael J Mlodzianoski, Hannah D Coughlan, Naiara G Bediaga, Gaetano Naselli, Erin C Lucas, Qike Wang, Carolyn A de Graaf, Douglas J Hilton, Leonard C Harrison, Gordon K Smyth, Kelly L Rogers, Thomas Boudier, Rhys S Allan, Timothy M Johanson

**Author notes:** Corresponding author: Timothy Johanson.

## Abstract

The proximity pattern and radial distribution of chromosome territories within spherical nuclei are well understood to be random and non-random, respectively. Whether this distribution pattern is conserved in the partitioned or lobed nuclei of polymorphonuclear cells is unclear. Here we use chromosome paint technology and a novel high-throughput imaging analysis pipeline to examine the chromosome territories of all 46 chromosomes in hundreds of single human neutrophils – an abundant and famously polymorphonuclear immune cell.

By comparing the distribution of chromosomes to randomly shuffled controls, and validating with orthogonal chromosome conformation capture technology, we show for the first time that all human chromosomes randomly distribute to neutrophil nuclear lobes, while maintaining a non-random radial distribution within these lobes. Furthermore, by leveraging the power of this vast dataset, we are able to reveal characteristics of chromosome territories not detected previously. For example, we demonstrate that chromosome length correlates with three-dimensional volume not only in neutrophils but other human immune cells.

This work demonstrates that chromosomes are largely passive passengers during the neutrophil lobing process, but are able to maintain their macro-level organisation within lobes. Furthermore, the random distribution of chromosomes to the naturally partitioned nuclear lobes suggests that specific transchromosomal interactions are unimportant in mature neutrophils.

## Introduction

First proposed in 1885 (1), interphase chromosomes maintaining a territorial organisation is now a widely accepted principle of nuclear organisation in most eukaryotes (2). This is unsurprising given the importance of this organisation to functions as fundamental as gene expression and DNA repair. For example, the radial position of a chromosome within the nucleus is strongly correlated with its transcriptional activity (3, 4). Furthermore, the proximity of chromosomes to one another (both homologous and non-homologous) is thought to be important during DNA repair (5) and potentially even in direct gene regulation (6-10).

While the radial distribution of chromosomes is well understood to be non-random (2), the position of chromosomes relative to each other, or proximity pattern, is contentious, with reports of both non-random (11-15) and random distributions (16). In the absence of being able to observe interphase chromosome movements in live cells over long periods of time, combined with the absence of physical barriers to restrict chromosome movement, it is possible that these studies simply differ in their detection of transient or infrequent interphase chromosomal interactions or movements.

Human neutrophils constitute approximately two thirds of the immune cells in human blood. They are readily identifiable by their polymorphonuclear nature with their nuclei being segmented into 2-6 lobes joined only by thin filaments of nucleoplasm (17). Here we exploit this natural nuclear segmentation to examine the importance of interphase chromosome distribution. We hypothesise that if interactions between chromosomes are biologically important these chromosomes would preferentially locate together into neutrophil nuclear lobes, enabling continuing interaction.

Using chromosome paint technology alongside novel high-throughput image analysis pipelines, we have examined all 46 human chromosomes in 240 single neutrophils. We reveal that while the radial distribution of chromosomes within neutrophil nuclear lobes is non-random, the distribution of chromosomes to lobes is in fact random, suggesting that gene regulatory interactions between specific chromosomes are highly unlikely to occur in human blood neutrophils.

## Results

### Novel analysis pipeline detects the position and characteristics of all human chromosomes in three-dimensions

Given the complexity of examining all 46 human chromosomes in hundreds of single, segmented neutrophil nuclei, we developed a bespoke image analysis pipeline to detect the position and three-dimensional characteristics of all chromosomes in images generated using chromosome paint.

In brief, each of the 22 autosome pairs and the X and Y chromosomes within fixed healthy male human blood neutrophil nuclei (Supp Fig 1A) is “painted” with a specific combination of fluorescent oligonucleotides within the chromosome paint mix (Fig 1A, B). The whole nucleus is then imaged and analysed using our analysis pipeline (Fig 1C). First, the intensity of each of the five channels in each individual image is normalised. A malleable grid with lines that flex to incorporate nearby voxels of similar channel intensity patterns is then applied to each image. This flexibility allows the grid to capture the highly variable three-dimensional shapes formed by chromosomes. Adjacent cubes of the grid that share channel intensities are then combined to create “objects”. Based upon the expected spectral combinations for each chromosome (Fig 1 B) these objects are then assigned as chromosomes. To avoid the common pitfall of arbitrary thresholding to define genuine signal from background (2), here every image has an individually determined threshold. This threshold is calculated by automated sequential testing of various thresholds to determine the value which maximises the number of objects with genuine chromosome channel combinations, while minimizing spectrally spurious objects (e.g. detected objects that have a channel combination differing from the 24 specific chromosome combinations).

**Figure 1:**
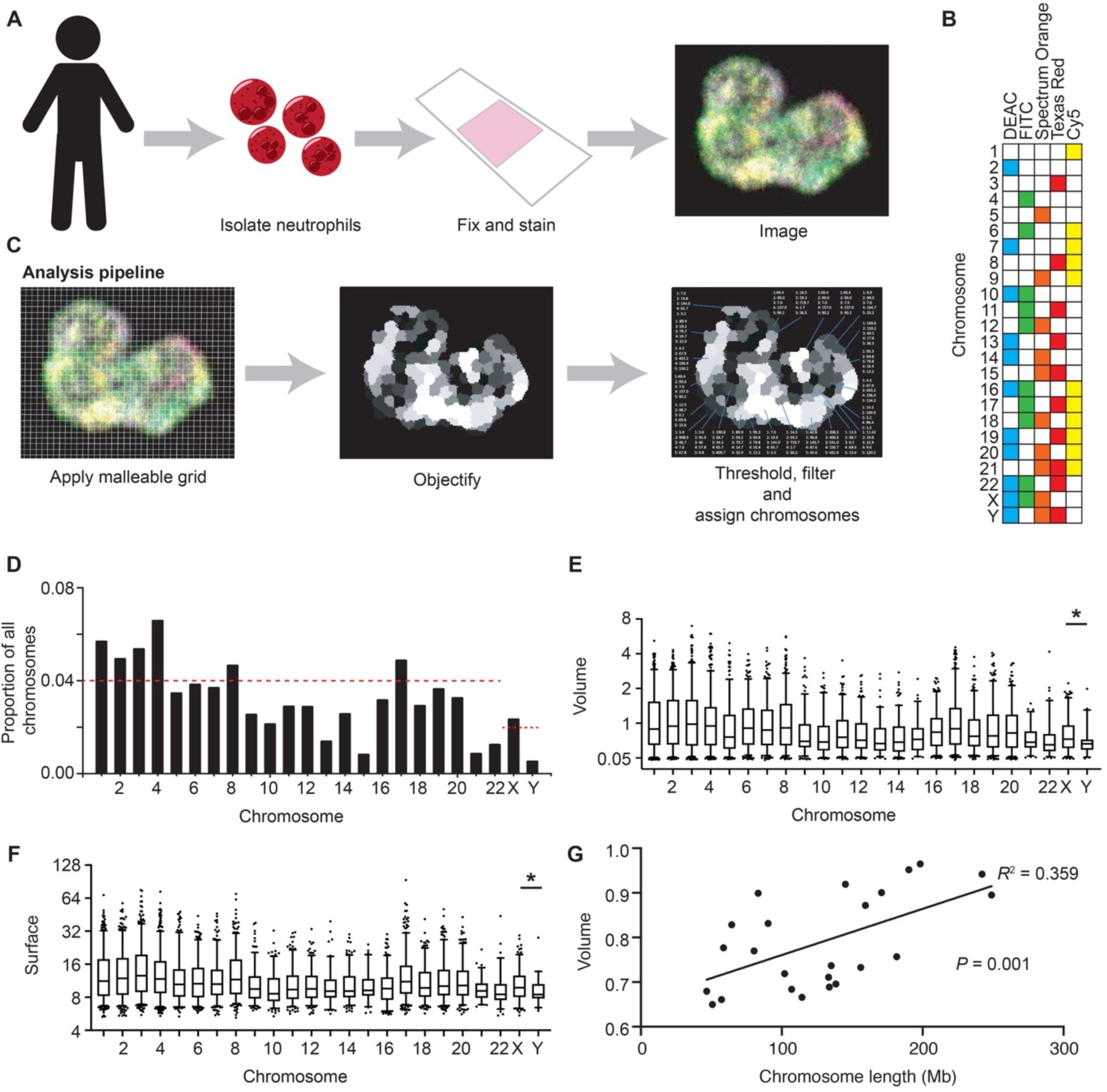
Novel analysis pipeline detects the position and characteristics of all human chromosomes in three-dimensions. (A-C) Schematic of neutrophil isolation, chromosome paint (A), spectral character of each human chromosome (B) and novel image analysis pipeline (C). (D) Proportion of total chromosomes detected made up by each chromosome in human blood neutrophils. Red line represents expected proportion if all 24 chromosomes were detected equally (0.04 for autosomes, 0.02 for sex chromosomes). (E, F) Box and whisker plot (5^th^-95^th^ percentile) of the volume (µm^3^) (E) and surface area (µm^2^)(F) of each chromosome across all 240 neutrophils. Unpaired T test used to compare X and Y chromosomes. (G) Plot of median chromosome volume across all neutrophils against chromosome linear length with straight line fitted (y=1.09e-3x+0.649, R^2^ = 0.359 and P = 0.001).

Importantly, chromosome paint and our image analysis pipeline detect the majority of the 22 human autosomes and the sex chromosomes at approximately the expected proportions (Fig 1 D, Supp Fig 1B). While some chromosomes appear more difficult to define or detect (e.g. chromosome 13, 15, 21 and 22), the proportions of chromosomes detected are similar across human immune cell types (CD4^+^ and CD8^+^ T cells) (Supp Fig 1C, D), suggesting that the variation in detection frequency observed is technical, not biological.

We next examine the three-dimensional character of all 46 chromosomes, including volume (Fig 1 E) and surface area (Fig 1 F), among others (Supp Fig 1E, F). While the physical characteristics of the chromosome territories varies greatly across neutrophil nuclei, the volume and surface area of the Y chromosomes are, as expected, consistently and significantly (P=0.02 and P= 0.04, respectively) smaller than the X chromosome (Fig 1 E, F).

While the differences in the three-dimensional character of the chromosomes are subtle, the ability to examine all chromosomes (as opposed to between 2-7 chromosomes (18-20)) in large numbers of cells affords the power to reveal correlations previously missed. For example, contrary to previous studies suggesting that there is no relationship between chromosome linear length and three-dimensional volume (18, 20), our analysis reveals a significant linear relationship between the two (r^2^ = 0.359, *P* = 0.001), not only in neutrophils (Fig 1G), but other human immune cells (Supp Fig 1G).

Thus, while chromosome paint data contains variance in both chromosome detection and three-dimensional parameters, the scale of our dataset allows elucidation of biologically important correlations and phenomena that were undetectable using previous methods.

### Human neutrophil chromosomes distribute randomly to nuclear lobes

There is conflicting evidence as to whether the proximity pattern of chromosomes is random or non-random, even within polymorphonuclear nuclei (18, 20-23).

In order to examine the distribution of chromosomes to neutrophil nuclear lobes, we first developed a high throughput pipeline to define the lobes. In brief, the pipeline uses watershed analysis from manually assigned seeds within each lobe (Supp Fig 2A) to rapidly define nuclear lobes (Supp Fig 2B). Consistent with previous studies (22), we find that the majority of healthy human blood neutrophils have between 3-4 lobes (Supp Fig 2C) of approximately equal volumes (Supp Fig 2D).

Next, we performed a lobe colocalization analysis to examine the frequency with which each chromosome is located within a neutrophil lobe with any other (Fig 2A). We find significant non-random distribution of 44 pairs of chromosomes (Supp Table 1). However, this analysis does not consider the character of the chromosomes. For example, a larger chromosome would likely have less lobe partners due simply to occupying more lobe volume. Thus, to avoid the confounding impacts of chromosome character, we repeated our colocalisation analysis on randomly shuffled controls (Fig 2B). To generate these shuffled images, we created an imaging framework that randomly places every chromosome (with its individual three-dimensional character and that of the associated nucleus preserved) within the nuclear space of each individual image. Chromosomes were not permitted to overlap one another or the nuclear boundary. This shuffling was performed five independent times for each of the 240 biological images. The colocalisation frequency was then examined in each set of shuffled images to determine the variability of shuffling, but more importantly the impact of chromosome character on lobe sharing frequency. The analysis revealed significant lobe sharing in the randomly shuffled images, suggesting that chromosome character does impact lobe distribution.

**Figure 2.**
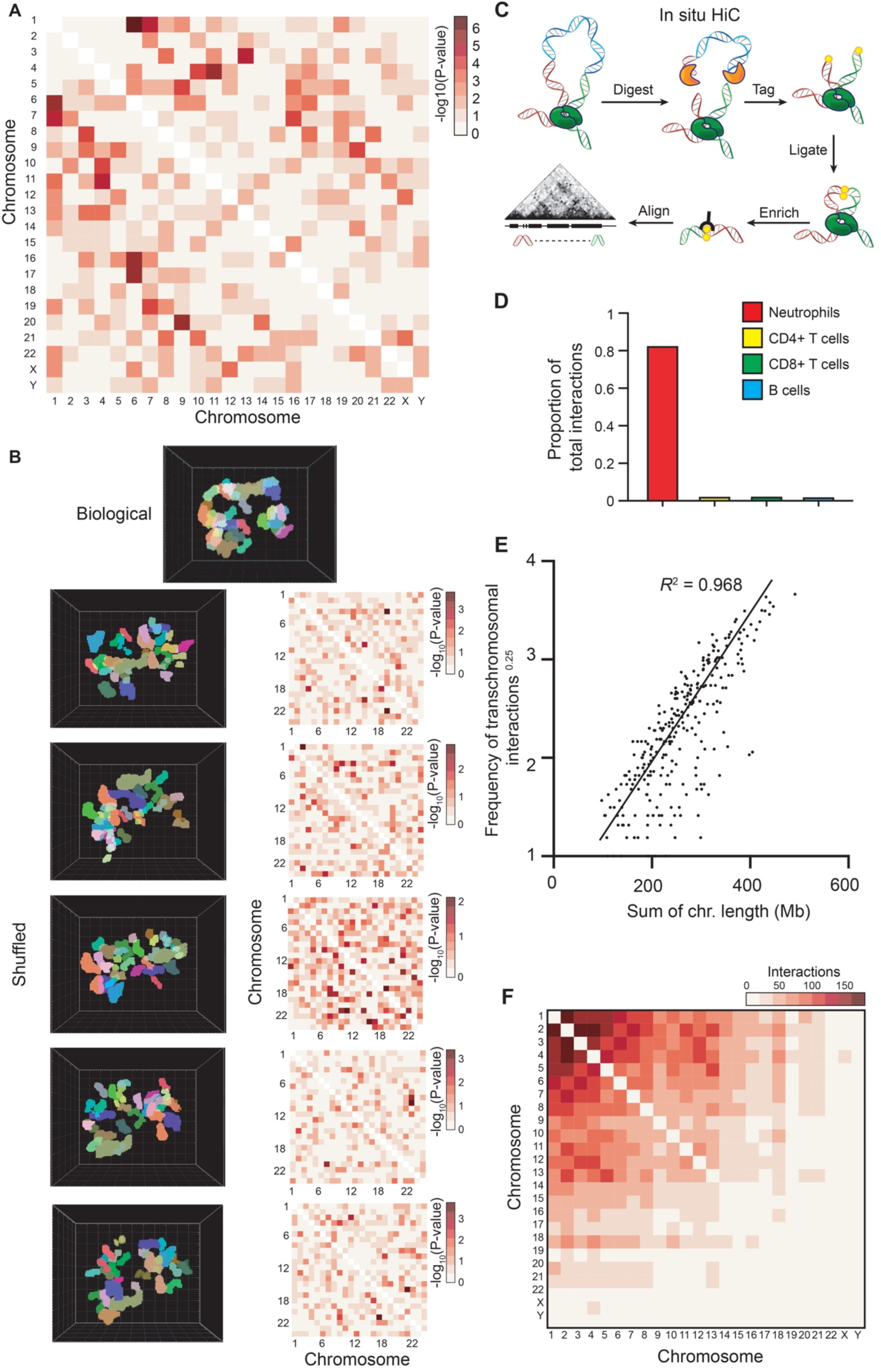
Human neutrophil chromosomes distribute randomly to nuclear lobes. (A) Heatmap of the -log_10_(p-value) from chromosome lobe colocalization analysis in human blood neutrophil nuclear lobes. The analysis determined if pairs of chromosomes are found colocalized within a lobe at a higher rate than expected by chance (B) Top: Three-dimensional render of chromosomes within a human neutrophil nucleus. Left: Three-dimensional renders of the same neutrophil and chromosomes after five sets of random chromosome shuffling. Right: Heatmap of the -log_10_(p-value) from the chromosome colocalization analysis of five independent sets of random chromosome shuffling. (C) Schematic of the proximity ligation reaction central to the *in situ* HiC protocol, (D) Proportion of total DNA-DNA interactions detected by *in situ* HiC and the *diffHiC* pipeline that occur between chromosomes (transchromosomal interactions) in human immune cells. (E) Frequency of transchromosomal interactions to the power of 0.25, plotted as a function of summed chromosome length in human neutrophils. Straight line fitted to data with no intercept (y=8.621e-3x, R^2^ = 0.9683 and p-value <2.2e-16). (F) Heatmap of the number of transchromosomal interactions by each chromosome to all others in human neutrophils.

To overcome the influence of chromosome character on lobe distribution, we filtered our significant lobe partner pairs (Supp Table 1) by significance of co-lobe localisation greater than that observed in the randomly shuffled analysis. Eleven chromosome pairings surpassed this threshold (Supp Table 2), suggesting that these chromosomes may preferentially co-localise in neutrophil nuclear lobes. However, the pairs in question share very similar spectral combinations, suggesting that perhaps the significance observed is an artefact of miscalling. For example, if one region of chromosome 1(painted Cy5) was occasionally miscalled as chromosome 6 (painted Cy5 and FITC) or vice versa this could lead to a significant colocalization score.

To validate our chromosome paint findings, we performed an orthogonal analysis of chromosome proximity using *in situ* HiC (24). *In situ* HiC utilises proximity ligation to determine the three-dimensional association of regions of DNA in populations of fixed nuclei (Fig 2C). While in most cells the vast majority of ligation events occur within chromosomes, transchromosomal interactions can be used as a measure of chromosome-chromosome proximity (24). Performing *in situ* HiC on human neutrophils we find that they exhibit dramatically more transchromosomal interaction than human immune cell types with spherical nuclei (Fig 2D). This is likely due to the increased physical confinement within neutrophil nuclei (25). If, as hinted at by chromosome paint, particular chromosomes are more frequently co-localised to neutrophil nuclear lobes than expected by chance, we would expect to observe an enrichment of transchromosomal interactions between these chromosomes compared to other chromosome pairs. However, we find that the distribution of transchromosomal interactions in human blood neutrophils is predominantly dependent on chromosome length (r^2^ = 0.968, Fig 2E) with no notable increase in interaction observed between any of the chromosome pairs suggested to co-localise by chromosome paint (Fig 2F).

Thus, using two independent methods, and by comparing to randomly shuffled controls, we find no consistent evidence of non-random chromosome distribution to human neutrophil nuclear lobes, suggesting that the distribution is in fact random.

### Human neutrophil chromosomes do not position randomly within nuclear lobes

Consistent with findings in spherical nuclei (16), our distribution analysis finds that the proximity pattern of chromosomes within neutrophil nuclei is random. Alongside random proximity pattern, non-random radial chromosome distribution is also well documented. As such, larger chromosomes are consistently observed closer to the nuclear periphery than their more diminutive counterparts (16). This has even been observed in a handful of neutrophil chromosomes (19).

To determine whether all human neutrophil chromosomes exhibit this relationship with the nuclear periphery, we examine the position of chromosomes relative to the lobed nuclear membrane. To do so we first break the radius of each lobe into a continuous scale from 0 to 1, 1 being the three-dimensional centre of the lobe and 0 being the nuclear periphery (Fig 3A). We then give each chromosome within the lobe a value between 0 and 1 based upon the mean position of its total volume (Fig 3B). By applying this method to all chromosomes in all lobes we are able to determine the position of each chromosome relative to the lobe boundary. As such, we find a significant relationship between chromosome linear length and its relative position within a lobe (r^2^ = 0.749, *P* = 4.60E-08), with larger chromosomes more likely to be found nearer the nuclear periphery than shorter chromosomes (Fig 3C). Importantly, when we apply this same method to our randomly shuffled control images the association is no longer detected (Fig 3D), strongly suggesting that it is not simply the volume of the larger chromosomes underlying the relationship.

**Figure 3.**
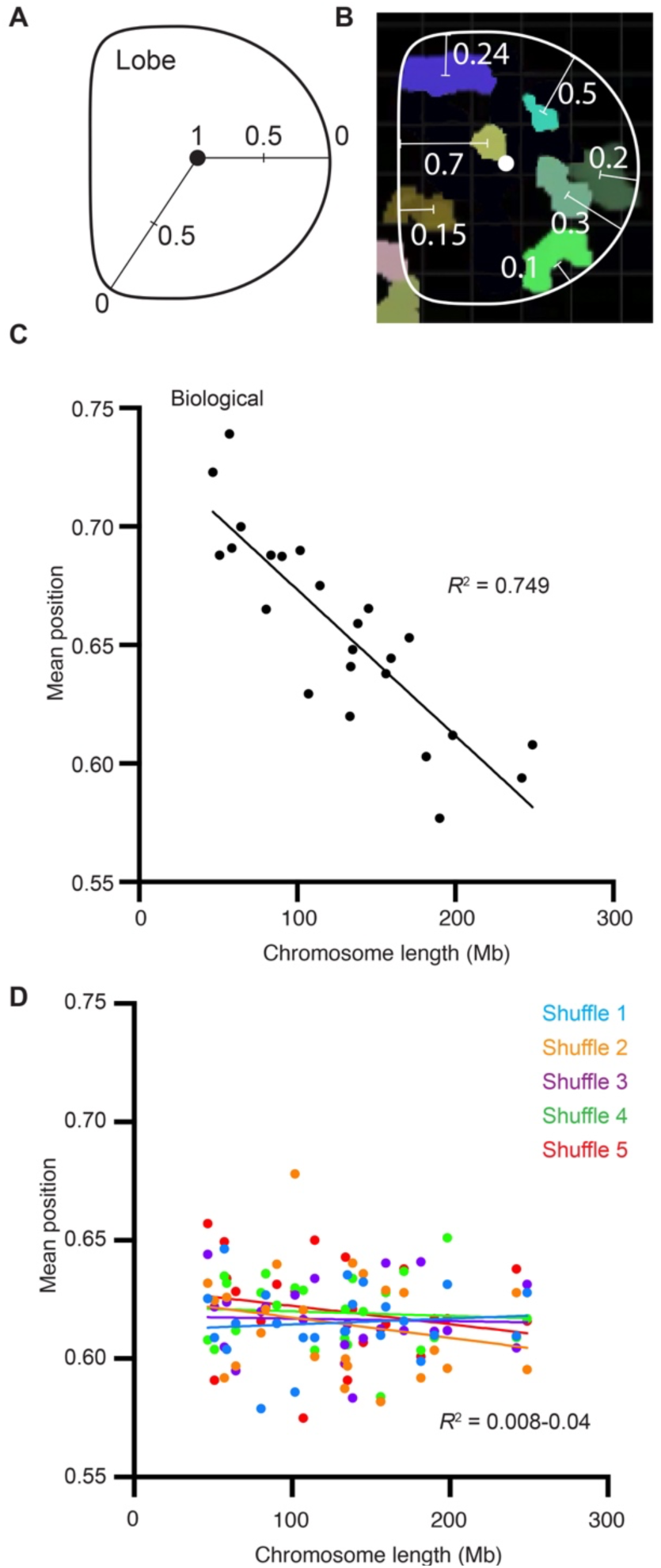
Human neutrophil chromosomes do not position randomly within nuclear lobes. (A) Schematic showing how radial position values within a neutrophil nuclear lobe are calculated (B) A slice through a neutrophil nuclear lobe image showing the approximate radial position values of the chromosomes. For clarity most chromosomes are not shown. (C) Scatterplot of the median of the mean chromosome volume radial position within human neutrophil nuclear lobes plotted as a function of chromosome length (Mb). Straight line fitted to data (y=-6.18e-4x+0.735, R^2^ = 0.7498 and p-value 4.6e-8). (D) Scatterplot of the median of the mean chromosome volume radial position from five independent sets of random chromosome position shufflings plotted as a function of chromosome length (Mb). Straight line fitted to each instance independently (y=-2.38e-5x+0.612, R^2^ = 0.008 and p-value = not significant (ns), y=-8.47e-5x+0.612, R^2^ = 0.0473 and p-value ns, y=-1.24e-5x+0.618, R^2^ = 0.0023 and p-value ns, y=-2.12e-5x+0.622, R^2^ = 0.0066 and p-value ns, y=-7.45e-5x+0.630, R^2^ = 0.0378 and p-value ns).

Thus, as in spherical nuclei, chromosomes within the highly physically restricted neutrophil nuclear lobes are positioned in a non-random and size-influenced manner.

## Discussion

Here we outline an automated method for the analysis of the three-dimensional character and position all 46 chromosomes in hundreds of single cells. By leveraging the power provided by this vast data set we resolve a number of longstanding questions in the field of chromosome organisation. We demonstrate in numerous immune cell types that chromosome length does in fact have a linear relationship with chromosome volume. Moreover, in human neutrophils we show that chromosome lobe distribution is random, while chromosome radial position within lobes is non-random.

The similarities in chromosome position between lobed neutrophil nuclei and spherical nuclei suggest that neutrophil chromosomes simply move and adapt to the more restricted nuclear environment created by lamin-mediated constriction (26). Thus, it appears that most chromosomes are simply passengers in the lobing process. However, there is one chromosome that appears to influence nuclear morphology – the inactive X chromosome. As such, the inactive X chromosome is frequently located within a small appendage protruding from the terminal lobe of female neutrophil nuclei (18, 22, 27). Strong evidence that the inactive X chromosome influences the formation of these appendages comes from the neutrophils of XXX or XXXX individuals which frequently exhibit two or three appendages, respectively (18). How the inactive X chromosome itself, its epigenetic state or its associated factors drive the formation of the nuclear appendages is unclear. However, these unique appendages do suggest that chromosomes are not always simply passive during nuclear morphology changes, but can under certain circumstances can influence the process.

Compared to their precursors and other immune cells, mature neutrophils are transcriptionally muted (19, 28, 29). It is unclear why. Prior to our study, one possibility was that neutrophil lobing perturbed the macroscale organisation of chromosomes, and thus their transcriptional activity. However, consistent with examinations of fewer (between 4-7) chromosomes (18, 19), we find that chromosome territories remain highly organised within neutrophil nuclear lobes. Thus, overt chromosomal disorganisation does not appear to explain the relative transcriptional inactivity of neutrophils.

It has been previously reported that regions on one chromosome can influence the expression of a gene on another, presumably via three-dimensional physical proximity (30). These interactions are known as gene regulatory transchromosomal interactions (31). Interestingly, these interactions have been predominantly reported in immune cells (6-10). However, both their function and indeed existence is debated (32). Here, while we reveal relatively large numbers of transchromosomal interactions in neutrophil nuclei via *in situ* HiC, our data demonstrate little to no preferential distribution of chromosomes to neutrophil nuclear lobes (where chromosomes are physically constrained and could physically interact only with their lobe mates), which suggests that gene regulatory transchromosomal interactions do not occur in mature human neutrophils.

The non-random radial distribution of chromosomes we have detected in neutrophil nuclear lobes is common in other cell types (16) and species (33). However, the cause and possible purpose of this organisation is not known. It has been proposed that the organisation could be due to the differential timing of centromere separation (34) or chromosome position during mitosis (16), differential interaction between chromosomes and the nuclear periphery and/or other nuclear bodies (35) or simply the force of transcription from highly transcribed chromosomes acting on those lowly transcribed (36). A number of purposes for the radial organisation have also been proposed, from protecting the genome from viruses and mutagens using a layer of highly repetitive and heterochromatic DNA (37) to providing nuclear stability or rigidity (16, 38). Given our findings in the highly malleable neutrophil nuclei the latter seems unlikely. While the function of peripheral heterochromatin is unclear in neutrophils or other cell types, there is one cell type in which the radial position of heterochromatin has a clear function. As the retinal cells of nocturnal animals develop, they invert the vast majority of their heterochromatin from its peripheral position to a central core (39). This remarkable transformation endows the mature retinal cell enhanced light channelling characteristics thus enhancing low light vision.

## Figure Legends

**Supp Fig 1.**
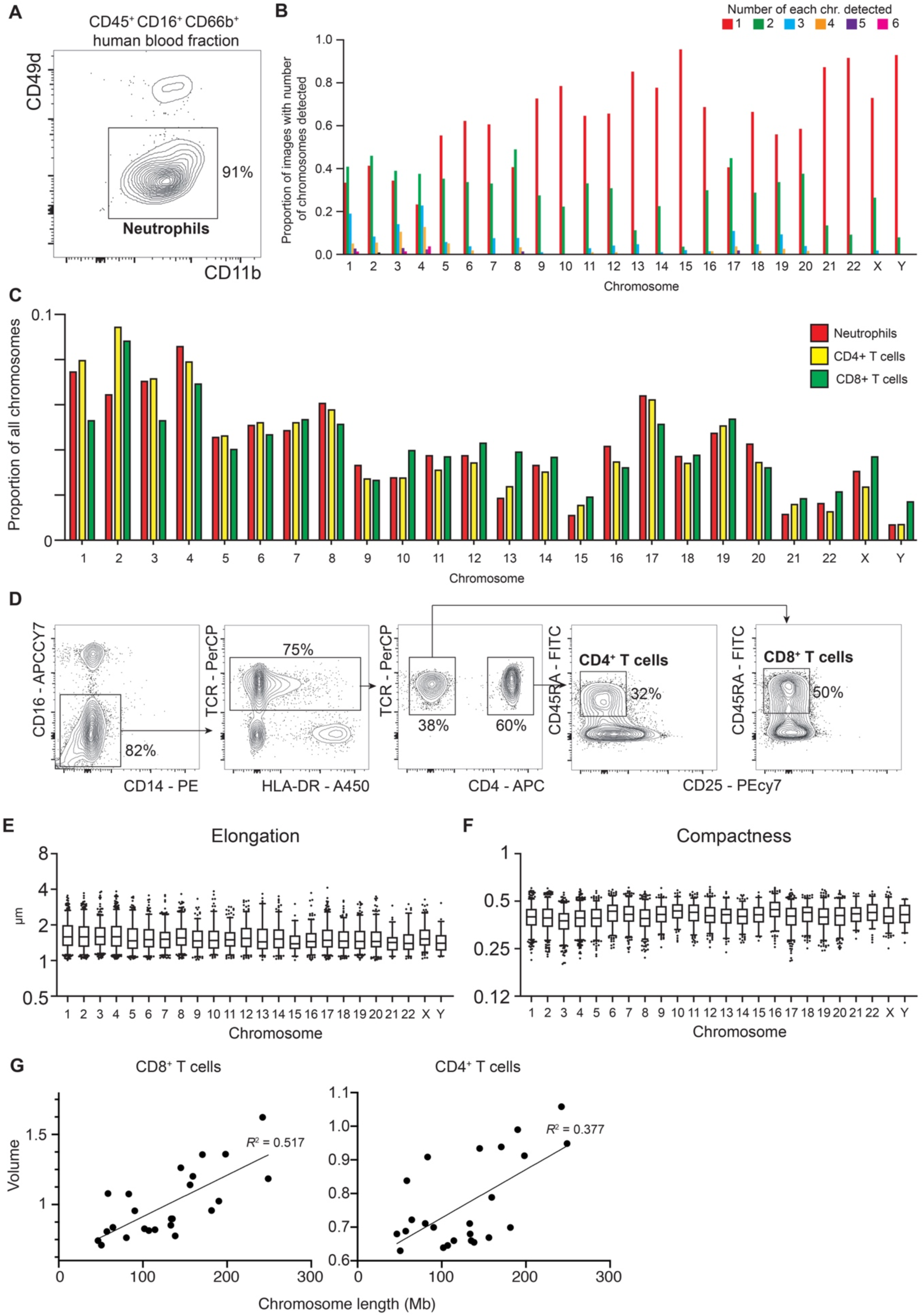
(A) Example purity check of enriched human blood neutrophils. (B) Number of chromosomes defined within a nucleus for all neutrophil images (proportion shown). (C) Proportion of total chromosomes detected made up by each chromosome in human blood neutrophils, CD4^+^ T cells and CD8^+^ T cells. Data from 171 CD4^+^ T cells and 30 CD8^+^ T cells. (D) Example sort profile of human blood CD4+ and CD8+ T cells. (E, F) Box and whisker plot (5^th^-95^th^ percentile) showing the elongation (E) and compactness (F) of chromosomes detected in human blood neutrophils. Elongation is the ratio between the largest axis of a fitted 3D ellipsoid and the second largest. Compactness is the normalized ratio between volume and surface. (G) Scatterplot of median chromosome volume in CD8^+^ T cells (left) and CD4^+^ T cells (right) against chromosome linear length. Data fitted with a straight line (y=2.94e-3x+0.621, R^2^ = 0.517 and p-value 7.5e-5 and y=1.41e-3x+0.588, R^2^ = 0.377 and p-value 0.00141, respectively).

**Supp Fig 2.**
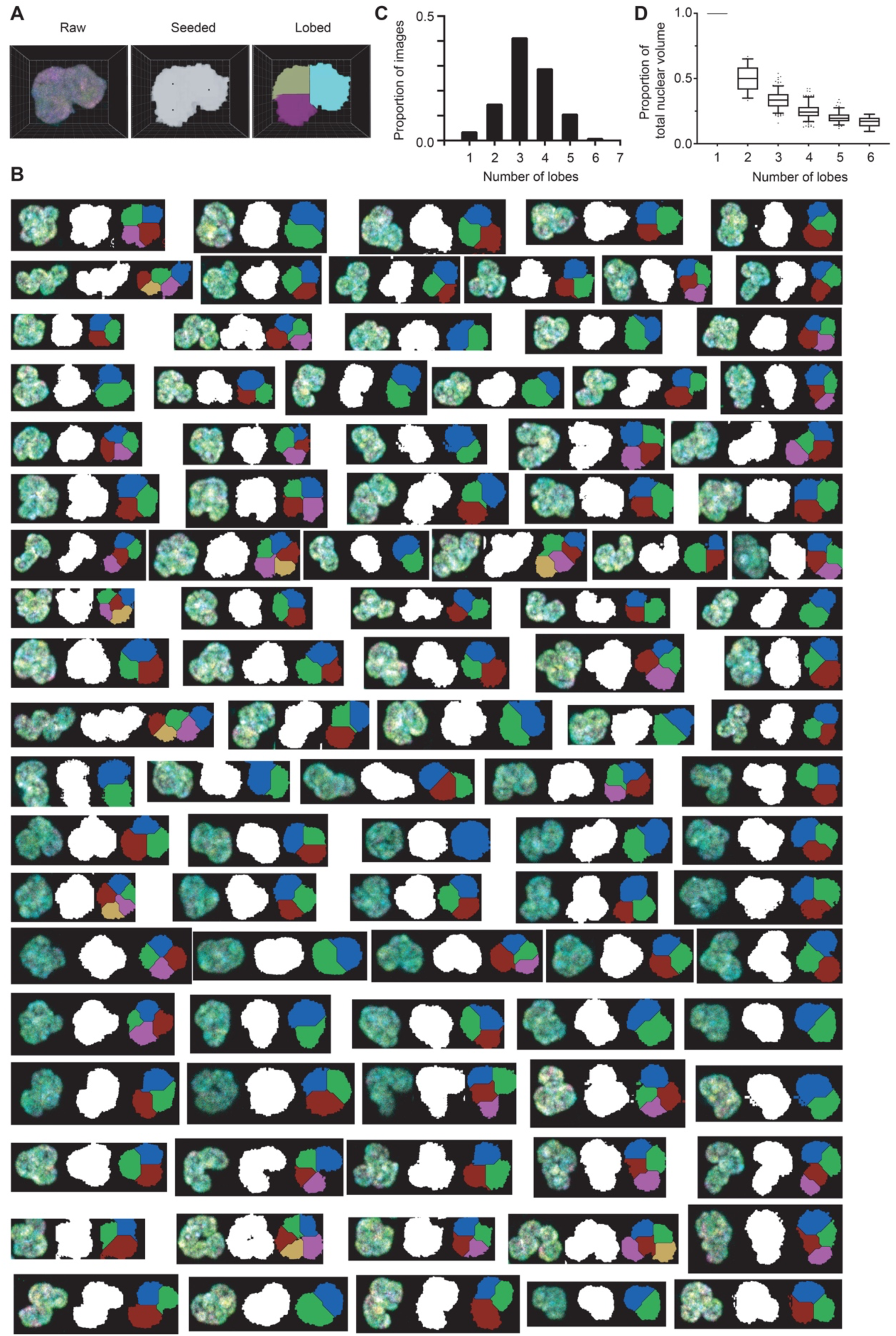
(A) Slice of a neutrophil image moving through the lobe calling pipeline, from raw image to the final lobed nucleus defined using watershed analysis. (B) Further examples of lobe calling pipeline results. (C) Bar plot showing the proportion of human blood neutrophils containing between 1-7 nuclear lobes. (D) Box and whisker plot (5^th^-95^th^ percentile) showing the volume of lobes as a proportion of the total nuclear volume in human blood neutrophils containing between 1-6 lobes.

## Methods

### Ethics Statement

Collection of human blood for research studies was approved by Human Research Ethics Committees of Melbourne Health and Walter and Eliza Hall Institute of Medical Research (application 88/03). Written consent was obtained.

### Cell isolation

Neutrophils were isolated from 2ml of healthy male donor whole blood following the EasySep Human Neutrophil Enrichment kit manufacturer’s protocol). Purity was ∼90%. Human T cells were isolated from the PBMC layer of the remaining whole blood after density separation in Leucosep™ tubes containing 15 mL Ficoll-Paque, following the manufacturer’s protocol. These cells were stained with TCRab-PerCP-eFluor 710 (eBioscience Cat.No. 46-9986-42), CD4-APC (BD Pharmingen Cat.No. 555349), CD25-PECy7 (BD Bioscience Cat.No. 557741), CD45RA-FITC (eBioscience Cat.No. 556626), CD14-PE (BioLegend Cat.No. 367104), CD16-APC-Cy7 (BD Bioscience Cat.No. 557758), HLA-DR-eFluor450 (eBioscience Cat.No. 48-9952-42) and CD19-BV650 (BioLegend Cat.No. 302238). CD4+ T cells (CD16-CD14-TCRab+ CD4+ CD45RA+ CD25-) and CD8+ T cells (CD16-CD14-TCRab+ CD4-CD45RA+ CD25-) were sorted to a purity >97%.

### Chromosome paint

Sixty thousand cells were settled on a poly-L-Lysine (Sigma Aldrich Cat no. P4707) coated cover slip at 37°C for 20 minutes, washed with phosphate buffer saline, before fixing with fresh 3:1 methanol:acetic acid (glacial) for 15 minutes at 22°C. The cells were then rinsed twice with water before being incubated in 2x saline-sodium citrate buffer (SSC) for 2 minutes. The cells were then dehydrated in an ethanol series (75%, 85% and 100%) for 2 minutes each. 7uL of Metasystems 24XCyte Human multicolour FISH probes (Metasystems Cat no. D-0125-060-DI) was pre-warmed on a glass slide at 37°C for 5 minutes before a further 2 minutes at 75°C with the cells added. The cells were then sealed and incubated at 37°C for 18 hours in a dark, humid chamber. After hybridisation was complete the cells were immersed in 72°C 0.4x SSC for 2 minutes, 2x SSC with 0.05% Tween-20 for 30 seconds before being sealed on a glass slide in 85% glycerol.

### Confocal Microscopy

Imaging experiments were performed on a Zeiss 880 confocal microscope using a 63x 1.4 NA objective lens. The system was run in lambda mode, recording fluorescence signal in 32 spectral channels over a spectral range of 410 nm to 690 nm (in 8.9 nm increments). Samples were imaged in 3D using Nyquist sampling of 70 nm pixel size and z-steps of 200 nm using the 405 nm, 488 nm, 561 nm and 633 nm lasers to excite the samples consisting of the fluorescent labels DEAC, FITC, Spectrum Orange, Texas Red and Cy5. Single colour control experiments were performed to determine the spectral signatures using the above microscope settings.

Tetraspeck beads (ThermoFisher) adhered to a coverslip and mounted to a microscope slide were imaged in 3D. Custom written Matlab (Mathworks, Natick, MA, USA) scripts measured the axial positions of the Tetraspeck beads in each spectral channel and removed the chromatic aberrations from the sample images. The five spectral signatures from the single colour controls were used to linearly unmix the 32-channel spectral fluorescence signal in each voxel into the five constituent fluorophores.

### Chromosome assignment and measurement

After channel unmixing and normalisation, voxels with similar values in the 5 channels are clustered together using a 3D Simple Linear Iterative Clustering (SLIC) algorithm (40). Thresholding is then applied to the 5 channels and chromosome identity is assigned according to known channel combinations. The threshold is set automatically by examining a range of thresholds and select that which yields the optimal number of chromosome objects (2) and the minimum number of false combination objects. Chromosome measurements are performed on assigned chromosomes using algorithms and tools from the 3D ImageJ suite (41). 3D Eroded Volume Fraction (42) is also performed to compute the position of chromosomes within the nucleus.

### Nuclear lobe calling

Nuclei boundaries are detected by summing all channels and global thresholding. The approximate 3D positions of the lobe centres are manually marked before a watershed method separates the nucleus into lobes.

### Chromosome shuffling

While considering the chromosomes three-dimensional character, the centre positions of all chromosomes are randomly distributed within the nuclear space until all chromosomes fit and no overlapping is observed between chromosomes. Due to space constraints shuffling was not possible for every nucleus attempted.

All images were stored within an OMERO Database (43), processing and analysis were then automated using the TAPAS home system (44).

### Chromosome measurements analysis

Chromosome measurements were analysed with R using packages dplyr and purr. Objects not assigned to a known combination of values were removed for analysis and plotting. Straight lines were fitted to the data with function lm() from R.

### Chromosome co-localisation in lobes

Nuclei with less than ten chromosomes detected or just one lobe were excluded from the analysis. For the remaining nuclei, the volume of each chromosome in each lobe was recorded. The co-localisation score of any two chromosomes in a nucleus was quantified by *Sum(p*_*il*_ *\* p*_*jl*_ ), where *p*_*il*_ is the proportion by volume of chromosome *i* in lobe *l, p*_*jl*_ is the proportion by volume of chromosome *j* in lobe *l*, and the sum is over all lobes in the nucleus. Co-localization p-values were obtained as follows. Each chromosome was treated in turn as the reference chromosome. The co-localisation scores for the reference chromosome with each other chromosome were ranked within each nucleus. The ranks were summed across nuclei and converted to z-scores assuming uniformly distributed ranks for each nucleus. The p-values were adjusted for multiple testing using the Bonferroni correction. Heatmaps of the -log_10_(p-value) were generated using the R package gplots with the function heatmap.2.

### *In situ* HiC

*In situ* HiC was performed as previously described (24) for one biological replicate of primary human neutrophils. The library was sequenced on an Illumina NextSeq 500 to produce 81-bp paired-end reads. HiC libraries for CD4^+^ T cells, CD8^+^ T cells and B cells are from GSE105776. The data pre-processing and analysis was performed with the *diffHic* pipeline (45) in R as previously described with changes in parameters (32). Where biological replicates were available, the libraries were summed after pre-processing.

Chromosomal looping interactions were detected using a method similar to that described previously by Rao et al. (24) and in (46). Read pairs were counted in bin pairs of 50 kbp anchors for all libraries. For each bin pair, the log-fold change over the average abundance of each of several neighbouring regions was computed. Neighbouring regions in the interaction space included a square quadrant of sides ‘x+1’ that was closest to the diagonal and contained the target bin pair in its corner; a horizontal stripe of length ‘2x+1’ centred on the target bin pair; a vertical stripe of ‘2x+1’, similarly centred; and a square of sides ‘2x+1’, also containing the target bin pair in the centre. The enrichment value for each bin pair was defined as the minimum of these log-fold changes, i.e., the bin pair had to have intensities higher than all neighbouring regions to obtain a large enrichment value. The neighbourhood counts for the libraries at 50 kbp bin size were computed with the neighborCounts function from the *diffHic* package with flank (x) 5 bin sizes (i.e., 250 kbp) and enrichment values were determined with filterPeaks function with get.enrich=TRUE. Looping interactions were then filtered with filterPeaks. Loops were defined as those with enrichment values above 1, were more than 100 kbp from the diagonal and with minimum count greater than 10 for all libraries except the neutrophils which used a minimum count of 5 to accounts for differences in library size. Directly adjacent loops in the interaction space were aggregated into clusters to a maximum cluster size of 500 kbp using the clusterPairs function from the *csaw* package v1.18.0 (47). Blacklisted genomic regions were obtained from ENCODE for hg38 (48). Loops that that had at least one anchor in a blacklisted genomic region were removed.

Heatmaps of the loops between chromosomes where generated using the R package gplots with the function heatmap.2. The frequency of transchromosomal interactions to the power of 0.25 was plotted as a function of the sum of the chromosome lengths. A linear model was fitted to data with the lm function.

## Data availability

Sequence data that support the findings of this study are tabulated in the supplementary tables and are available in the GEO database under accession number GSE157229.

## Acknowledgments

The authors thank S. Johanson for constructive discussions, and the volunteers at the Walter and Eliza Hall Volunteer Blood Donor Registry for their donation.

